# Unexpected arabinosylation after humanization of plant protein *N*-glycosylation

**DOI:** 10.1101/2021.12.17.473187

**Authors:** Lennard L. Bohlender, Juliana Parsons, Sebastian N. W. Hoernstein, Nina Bangert, Fernando Rodríguez-Jahnke, Ralf Reski, Eva L. Decker

**Author notes:** **Corresponding author:** Eva L. Decker, Plant Biotechnology, Faculty of Biology, University of Freiburg, Freiburg, Schaenzlestrasse 1, 79104 Freiburg, Germany.

## Abstract

As biopharmaceuticals, recombinant proteins have become indispensable tools in medicine. An increasing demand, not only in quantity but also in diversity, drives the constant development and improvement of production platforms. The *N*-glycosylation pattern on biopharmaceuticals plays an important role in activity, serum half-life and immunogenicity. Therefore, production platforms with tailored protein *N*-glycosylation are of great interest. Plant-based systems have already demonstrated their potential to produce pharmaceutically relevant recombinant proteins, although their *N*-glycan patterns differ from those in humans. Plants have shown great plasticity towards the manipulation of their glycosylation machinery, and some have already been glyco-engineered in order to avoid the attachment of plant-typical, putatively immunogenic sugar residues. This resulted in complex-type *N*-glycans with a core structure identical to the human one. Compared to humans, plants lack the ability to elongate these *N*-glycans with β1,4-linked galactoses and terminal sialic acids. However, these modifications, which require the activity of several mammalian enzymes, have already been achieved for *Nicotiana benthamiana* and the moss Physcomitrella. Here, we present the first step towards sialylation of recombinant glycoproteins in Physcomitrella, human β1,4-linked terminal *N*-glycan galactosylation, which was achieved by the introduction of a chimeric β1,4-galactosyltransferase (FTGT). This chimeric enzyme consists of the moss α1,4-fucosyltransferase transmembrane domain, fused to the catalytic domain of the human β1,4-galactosyltransferase. Stable FTGT expression led to the desired β1,4-galactosylation. However, additional pentoses of unknown identity were also observed. The nature of these pentoses was subsequently determined by Western blot and enzymatic digestion followed by mass spectrometric analysis and resulted in their identification as α-linked arabinoses. Since a pentosylation of β1,4-galactosylated *N*-glycans was reported earlier, e.g. on recombinant human erythropoietin produced in glyco-engineered *Nicotiana tabacum*, this phenomenon is of a more general importance for plant-based production platforms. Arabinoses, which are absent in humans, may prevent the full humanization of plant-derived products. Therefore, the identification of these pentoses as arabinoses is important as it creates the basis for their abolishment to ensure the production of safe biopharmaceuticals in plant-based systems.

## 1 Introduction

Recombinant protein biopharmaceuticals are highly effective and specific, and therefore essential in the area of healthcare. The advancement of biotechnology made their production feasible, their share in the market has grown steadily in the last decades and is predicted to keep growing (Facts and Figures 2021: The Pharmaceutical Industry and Global Health; Walsh, 2018). The production of high-quality therapeutic proteins is still a complex process. For this the biosynthesis machinery from cells is required, and the choice of the production platform is highly associated with the product’s requirements and quality (Tripathi and Shrivastava, 2019). Proteins are frequently post-translationally modified. Particularly, protein *N*-glycosylation, a very common post-translational modification (PTM) in most eukaryotes, is of great importance as most protein biopharmaceuticals need a correct glycosylation to achieve the desired therapeutic efficacy (Solá and Griebenow, 2010) and to prevent immunogenic effects by the pharmaceutical (Zhou and Qiu, 2019). Mammalian (esp. Chinese Hamster Ovary (CHO)) cell lines, have dominated the recombinant biologics industry since the 1990s, largely because their PTMs resemble human ones (Walsh, 2018; Tripathi and Shrivastava, 2019). However, high production costs of these systems and the increasing demand of newly designed protein therapeutics, driven by the growing knowledge of molecular mechanisms of diseases, reveal the need for alternative platforms for tailored production. The current COVID-19 pandemic highlights particularly the urgent need to expand the production capacities for vaccines, diagnostic reagents and therapeutical proteins, such as neutralizing antibodies. Plant-based production of biopharmaceuticals offers an interesting alternative. For this, plants combine several advantageous properties like their ability to produce, fold and post-translationally modify complex proteins, a high range of scalability combined with cost-effective cultivation and the lack of human pathogens which provides inherent safe products (Buyel, 2019). Currently, one plant-produced recombinant therapeutic is on the market (Elelyso®, a β-glucocerebrosidase for the treatment of Morbus Gaucher, Grabowski et al., 2014) and many promising plant-made biopharmaceuticals are in clinical trials. Among them are the HIV-neutralizing human monoclonal antibody 2G12 produced in *Nicotiana tabacum* (Ma et al., 2015), the *Nicotiana benthamiana*-derived virus-like particles as candidate vaccines against influenza, dengue fever or COVID-19, respectively (Ward et al., 2020, 2021; Ponndorf et al., 2021) or α-galactosidase for enzyme replacement therapy in Morbus Fabry treatment produced in the moss Physcomitrella (Shen et al., 2016; Hennermann et al., 2019). These promising candidates demonstrate the potential of plant-based systems in this field. All these approved or in advanced clinical trials plant-derived biopharmaceuticals have in common, that their efficacy is not impaired by the lack of mammalian-typical *N*-glycosylation patterns, which differ from those produced in plants. The early processing of *N*-glycans in plants and mammals is conserved, while their maturation in the Golgi apparatus differs (Gomord et al., 2010). Plant and human *N*-glycans share the identical heptasaccharide GlcNAc_2_Man_3_GlcNAc_2_ (GnGn) di-antennary complex-type core structure, while fucosylation of the Asn-linked N-acetylglucosamine (GlcNAc) is α1,3-linked in plants and α1,6-linked in humans. In humans though not in plants, the GnGn core is extended via β1,4-linked galactose, which is often terminally capped with α2,6-linked sialic acid. In plants, the GnGn core is substituted with a β1,2-linked xylose, a sugar not produced in humans, and it is terminally extended by β1,3-linked galactose and α1,4-linked fucose, both linked to the outer GlcNAc residues, forming the trisaccharide Lewis A (Le^a^) epitope. This epitope as well as the plant-specific β1,2-attached xylose and the α1,3-attached fucose have been associated with antibody formation in humans (Fitchette et al., 1999; Wilson et al., 2001). Antibodies recognizing a therapeutic protein can affect its efficacy by altering the pharmacokinetics and pharmacodynamics, and represent an additional safety risk (Tourdot and Hickling, 2019). Therefore, to avoid potential immunogenicity of plant-made therapeutical proteins, plant-specific *N*-glycan residues have already been tackled. Plant-specific *N*-glycan xylosylation and fucosylation were eliminated in several plant-based systems by knockout (KO) or downregulation of the genes encoding the respective xylosyltransferases (XT) and fucosyltransferases (FT) (Koprivova et al., 2004; Strasser et al., 2004, 2008; Cox et al., 2006; Sourrouille et al., 2008; Shin et al., 2011; Hanania et al., 2017; Mercx et al., 2017; Jansing et al., 2018). Additionally, Le^a^ epitope formation was abolished in Physcomitrella by knockout of the β1,3-galactosyltransferase 1 (GalT1) encoding gene (Parsons et al., 2012). The triple KO of *xt, ft* and *galt1* in Physcomitrella resulted in an outstanding *N*-glycan homogeneity, with a strongly predominant GnGn glycosylation pattern (Parsons et al., 2012). This provides a suitable platform for the further glyco-optimization, comprising β1,4-galactosylation and sialylation.

The impact of terminal *N*-glycan residues on efficacy and functional role of protein therapeutics has been extensively reviewed (Jefferis, 2009; Li and d’Anjou, 2009; Solá and Griebenow, 2010). Terminal *N*-glycan sialylation increases the protein surface charge and hides the underlying sugars galactose, GlcNAc and mannose. Renal filtration and elimination rates are retarded for highly charged proteins (Solá and Griebenow, 2010). Additionally, liver asialoglycoprotein receptors recognizing terminal galactose, as well as mannose receptors, mainly on immune cells, recognizing terminal mannose or GlcNAc, are responsible for a rapid clearance of non-sialylated glycoproteins from serum (Datta-Mannan, 2019).

To reach *N*-glycan sialylation, which has already been stably attained in *N. benthamiana* and Physcomitrella (Kallolimath et al., 2016; Bohlender et al., 2020), the galactosylated *N*-glycan acceptor should be provided as a first step. *In planta N*-glycan β1,4-galactosylation has been achieved via expression of heterologous coding sequences (CDSs) of different versions of β1,4-galactosyltransferases (β1,4-GalT), including the sequences of various animal species along with the human one and chimeric varieties thereof (Palacpac et al., 1999; Bakker et al., 2001, 2006; Misaki et al., 2003; Huether et al., 2005; Fujiyama et al., 2007; Hesselink et al., 2014; Kittur et al., 2020; Kriechbaum et al., 2020). In these various approaches in different plant species, it has become evident that galactosylation efficiency and quality is influenced by diverse factors. Among them, localization of the enzyme within the Golgi apparatus plays an important role. When localized too early in the Golgi sub-compartments, the β1,4-GalT activity interferes with the activities of the α-mannosidase II (GMII) or the N-acetylglucosaminyltransferase II (GnTII), impeding further *N*-glycan maturation and leading to incompletely processed mono-antennary galactosylated *N*-glycans (Strasser et al., 2009; Schneider et al., 2015; Kallolimath et al., 2018). The localization of a protein anchored in the endomembrane system is dependent on the N-terminal cytoplasmic, transmembrane and stem (CTS) domain (Czlapinski and Bertozzi, 2006; Schoberer and Strasser, 2011). Accordingly, the CTS of the human β1,4-GalT, which is apparently localized in the early to medial plant Golgi apparatus was replaced by CTS sequences with an assumed late trans-Golgi localization. Chimeric variants of the β1,4-GalT with different CTS domains, like the CTS of the human sialyltransferase (Strasser et al., 2009), the CTS of the Arabidopsis β1,3-galactosyltransferase 1 (Kriechbaum et al., 2020) or the CTS of the Physcomitrella α1,4-fucosyltransferase (FTGT) (Bohlender et al., 2020) have been described and led to higher shares of di-antennary galactosylated *N*-glycans. Furthermore, the target glycoprotein itself influences its galactosylation efficiency (Kriechbaum et al., 2020), probably based on conformation-related accessibility.

In this study we analyzed the galactosylation efficiency of the chimeric β1,4-galactosyltransferase FTGT, which consists of the CTS domain of the moss α1,4-fucosyltransferase fused to the catalytic domain of the human β1,4-GalT (Bohlender et al., 2020), in Physcomitrella expressing human erythropoietin (hEPO) (Weise et al., 2007). Human EPO is a highly glycosylated protein hormone which inhibits apoptosis of erythroid progenitor cells and stimulates their differentiation, increasing the number of circulating mature red blood cells (Jelkmann, 2013). Recombinant hEPO (rhEPO) is widely used for the treatment of severe chronic anemia especially associated with chronic kidney disease and chemotherapy (Jelkmann, 2013). Additionally, non-sialylated rhEPO (asialo-rhEPO) is of pharmacological interest due to its tissue-protective activity devoid of erythropoietic activity (Peng et al., 2020).

FTGT expression led to a galactosylation efficiency of about 66% on rhEPO *N*-glycans, and 65% of the galactosylated fraction consisted of mature di-antennary galactosylated structures. However, up to five additional pentoses were found to be attached to about 92% of all β1,4-galactosylated *N*-glycans. Pentosylation on β1,4-galactosylated *N*-glycans was recently reported in *N. tabacum* and Physcomitrella (Bohlender et al., 2020; Kittur et al., 2020), indicating that this modification might affect different plant-based production systems; but so far no reports are available elucidating its identity. Here, we identified the unknown pentoses as α-linked arabinoses. The arabinose identity was verified by immunoblot-based detection on rhEPO with an anti-α1,5-arabinan antibody and specific digestion of the pentoses from rhEPO with α-L-arabinofuranosidase, confirmed via immunoblot and mass spectrometry analysis.

Arabinoses are not present in humans, and therefore potentially immunogenic (Anderson et al., 1984; Steffan et al., 1995; Leonard et al., 2005). Moreover, they might interfere with the efficient establishment of *in planta* sialylation. In this regard, the characterization of the undesired pentosylation as α-L-arabinosylation is an indispensable step towards the identification of the responsible glycosyltransferase and thus to provide plant-based glyco-engineered biopharmaceuticals with tailored *N*-glycosylation patterns.

## 2 Materials and Methods

### 2.1 Plant material and generation of transgenic moss lines

Physcomitrella (*Physcomitrium patens*) was cultivated as described previously (Frank et al., 2005). The recombinant human EPO (rhEPO)-producing moss line 174.16 (Weise et al., 2007; Parsons et al., 2013) is based on the Physcomitrella Δxt/Δft double knockout line lacking β1,2-xylosyltransferase and α1,3-fucosyltransferase activity (Koprivova et al., 2004, IMSC no.: 40828).This line produces and secretes rhEPO with a predominant GnGn-glycosylation pattern, partially decorated with additional Le^a^ epitopes (β1,3-galactosylation and α1,3-fucosylation) to the culture medium. The moss line Δ*galt1* was obtained previously by targeted knockout of the moss-endogenous β1,3-galactosyltransferase 1 (GalT1, Pp3c22_470V3.1) in line 174.16 (Parsons et al. 2012).

This line produces rhEPO devoid of any plant-specific sugar residues. Human-like β1,4-galactosylation was established based on the line 174.16 via the homologous integration of a chimeric β1,4-GalT-containing expression cassette (Bohlender et al., 2020) into the GalT1-encoding locus to achieve simultaneous GalT1 depletion. This chimeric variant, FTGT, contains the CTS domain of the moss-endogenous α1,4-fucosyltransferase (Pp3c18_90V3.1) fused to the catalytic domain of the human β1,4-GalT (NM_001497.4) and is driven by the long 35S promoter (Horstmann et al., 2004). Resistance to Zeocin was used to select transformed plants (Bohlender et al., 2020).

### 2.2 Protein precipitation from culture supernatant

For rhEPO production, the respective Physcomitrella lines were inoculated at an initial density of 0.6 g dry weight (DW) /L and cultivated for 10 days (Parsons et al., 2012). Recombinant hEPO was recovered from culture supernatant by precipitation with trichlorocetic acid as described before (Büttner-Mainik et al., 2011).

### 2.3 Enzymatic arabinose digestion

Protein pellets recovered from culture supernatant and containing moss-produced rhEPO were dissolved in a 100 mM sodium acetate buffer containing 2% SDS (pH 4.0). After 10 minutes shaking (1,200 rpm, Thermomix, Eppendorf) at 90°C and additional 10 minutes centrifugation at 15,000 rpm the supernatant was transferred to a fresh 1.5 ml reaction tube. Total protein concentration was determined using the bicinchoninic acid assay (BCA Protein Assay Kit; Thermo Fisher Scientific) following the manufacturer’s instructions. For each analyzed line, 10 μg of total protein were mixed with one unit of α-L-arabinofuranosidase from either *Aspergillus niger* or a corresponding recombinant version (E-AFASE or E-ABFCJ, Megazyme, Bray, Ireland) and incubated over night at 40°C. In parallel, enzyme-free samples from each moss line were treated under the same conditions.

### 2.4 SDS-PAGE and Western blot

For SDS-PAGE, samples of 5-10 μg protein were reduced with 50 mM dithiothreitol (DTT) for 15 minutes at 90°C and mixed with 4× sample loading buffer (Bio-Rad, Munich, Germany). Protein separation was carried out via SDS-PAGE in 12% polyacrylamide gels (Mini-PROTEAN® TGX™ Precast Gels, Bio-Rad, Munich, Germany) in TGS buffer (Bio-Rad) at 120 V. For molecular weight comparison the PageRuler™ Prestained Protein Ladder (26616, Thermo Fisher Scientific) was used. After electrophoretic separation, proteins were transferred to a polyvinylidene fluoride (PVDF) membrane (Cytiva) using a Trans-Blot SD Semi-Dry Electrophoretic Cell (Bio-Rad) with 1.5 mA / cm^2^ membrane for 1 h. After blotting, the membrane was blocked in 0.1% Tween20 in TBS (TBST) with 4% ECL Blocking Agent (Cytiva) at 4°C over night. For arabinose Western blots, the membrane was incubated for one hour at room temperature with LM6-M anti 1,5-α-L-arabinan antibody (Plant Probes, Leeds, UK) diluted 1:10 in TBST with 2% ECL Blocking Agent. After three times washing with TBST for 15 minutes, the blot was incubated with a peroxidase-linked rabbit anti-rat secondary antibody (Ab6250, Abcam, Cambridge, UK) diluted 1:25,000 in TBST with 2% ECL Blocking Agent. Detection was performed by chemiluminescence development (ECL™ Advance Western Blotting Detection Kit, Cytiva) according to the manufacturer’s instructions. For rhEPO Western blot the membrane was stripped after the arabinose Western blot. For this, the membrane was incubated two times for 10 minutes in mild stripping buffer (1.5% glycine (w/w), 0.1% SDS (w/w), 1% Tween20 (v/v), pH 2.2) and afterwards washed three times for 10 minutes with TBST under gentle shaking. After overnight membrane blocking (4% ECL Blocking Agent in TBST), anti-hEPO monoclonal antibody (MAB2871; R&D Systems, Minneapolis, USA) and peroxidase-linked anti-mouse secondary antibody (NA 9310V, Cytiva) in 1:4,000 and 1:100,000 dilutions, respectively, were used.

### 2.5 Mass spectrometry

The *N*-glycosylation pattern on rhEPO was analyzed via mass spectrometry (MS) on glycopeptides obtained by double digestion with trypsin and GluC. For this, the samples were reduced as described above and additionally S-alkylated with a final concentration of 120 mM iodoacetamide (IAA) for 20 minutes at RT in darkness prior to SDS-PAGE. After Coomassie staining as described previously (Bohlender et al., 2020), bands corresponding to the molecular weight of rhEPO, ranging between 20 kDa and 40 kDa, were cut. Double digestions were performed with trypsin (Promega. Walldorf, Germany) and GluC (Thermo Fisher Scientific) in 100 mM ammonium bicarbonate solution at 37°C overnight. Peptide recovery and sample cleanup were performed as described in Top et al. (2019). The initial MS analysis comparing the three test lines (I10, X13 and X24) was performed on an Q-TOF istrument as described in Michelfelder et al. (2017). Glycopeptides were identified from processed Mascot mgf files using custom Perl skripts. A precursor mass tolerance of 15 ppm was used to search for glycopeptide precursors. Glycopeptide identity was confirmed by the presence of typical *N*-glycan fragments such as GlcNAc oxonium ions (m/z-values: [GlcNAc]^+^ = 204.087, [GlcNAc - H_2_O]^+^ = 186.076, [GlcNAc - 2H_2_O]^+^ = 168.066, [GlcNAc - C_2_H_4_O_2_]^+^ = 144.065, [GlcNAc - CH_6_O_3_]^+^ = 138.055, [GlcNAc - C_2_H_6_O_3_]^+^ = 126.055) and glycan fragment ions ([GlcNAcHex]^+^ = 366.139, [GlcNAcHex_2_]^+^ = 528.191, [GlcNAcHexPent]^+^ = 498.1818, [GlcNAcHexPent_2_]^+^ = 630.2241, [GlcNAcHexPent_3_]^+^ = 762.266, [GlcNAcHexPent_2_]^+^ = 894.3087) at a fragment mass tolerance of 0.05 Da. Spectra were plotted from mgf files and labeled using custom Perl scripts an R (www.R-project.org). Lists of all searched precursor masses and identified glycopeptides are available in supplementary tables S1 and S2, respectively.

Further MS analyses were performed on a QExative Plus instrument (Thermo Scientific™, Bremen, Germany) as described previously (Top et al., 2019). Identification of glycopeptides and quantitation was performed as described in Bohlender et al. (2020). In brief, glycopeptides were identified with custom Perl scripts from Mascot mgf files of processed raw data. Precursors were matched at a mass tolerance of 5 ppm and resulting spectra were scanned for the presence of typical glycosylation reporter ions such as GlcNAc oxonium ions (m/z-values: [GlcNAc]^+^ = 204.087, [GlcNAc - H2O]^+^ = 186.076, [GlcNAc - 2H_2_O]^+^ = 168.066, [GlcNAc - C_2_H_4_O_2_]^+^ = 144.065, [GlcNAc - CH_6_O_3_]^+^ = 138.055, [GlcNAc - C_2_H_6_O_3_]^+^ = 126.055) and glycan fragment ions ([GlcNAcHex]^+^ = 366.139, [GlcNAcHex_2_]^+^ = 528.191. For fragment ion matching a mass tolerance of 0.02 Da was specified. Quantitative values (peak areas) for the identified glycopeptides were extracted using a custom Perl script from the allPeptides.txt file obtained from a default MaxQuant (V1.6.0.16, Cox and Mann (2008)) search on the respective raw data. A list of searched glycopeptides and their calculated precursor masses as well as a list of identified glycopeptides is available from supplementary tables S3 and S4. Quantitative values were normalized against the total intensity (sum of all peak areas) of the measurement and added together for precursors identified in different charge stages (table S5 and S6).

For a better visualization and representation of the results, the nomenclature for isomeric structures was simplified (e.g., AM, MA or a mixture of both are all displayed as AM). The presence of pentoses on known *N*-glycan structures is displayed as +P, while the range of detected pentoses on the corresponding structure is given in subscripted numbers, e.g. AM-structures with one to three attached pentoses are designated as AM+P_1-3_. A detailed breakdown of all identified structures and modifications is given in the supplementary table S4 and the quantification is depicted in the supplementary figure S2.

All mass-spectrometry data have been deposited to the ProteomeXchange Consortium via the PRIDE partner repository (Perez-Riverol et al., 2019) with the dataset identifier PXD030443.

## 3 Results

### 3.1 Expression of FTGT leads to efficient *N*-glycan β1,4 galactosylation on rhEPO and attachments of additional unknown pentoses

To achieve mature β1,4-galactosylation on rhEPO in moss, the plant 174.16, which produces rhEPO devoid of plant-specific xylose and α1,3-attached core fucose (Weise et al., 2007) was transformed with the expression construct coding for the chimeric β1,4-galactosyltransferase FTGT (Bohlender et al., 2020). This construct is targeted to the genomic locus encoding the β1,3-galactosyltransferase 1, *galt1*. Gene knockout via targeted integration in the *galt1* locus was confirmed by PCR, therefore presence of galactose on rhEPO glycopeptides can be inferred to be β1,4-linked and not β1,3 (Parsons et al., 2012). Three lines (I10, X13 and X24) were chosen for MS-based rhEPO glycopeptide analysis. A first MS survey revealed galactosylation in all three lines and on all three rhEPO *N*-glycosylation sites. However, in line I10 almost no di-antennary galactosylated structures were detected, and in line X13 a larger proportion of immature *N*-glycans, such as AM structures and a broader heterogeneity of *N*-glycans at the three different glycosylations sites, compared to line X24, were observed (Supplementary Table S2). Therefore, line X24 was chosen for further studies.

The X24-derived rhEPO glycopeptides displayed an average galactosylation efficiency of 66%, consisting of 43%di-antennary and 23% mono-antennary galactosylated structures. In addition to the expected galactosylation, single or multiple mass additions of 132.0423 Da, unknown from the *N*-glycans before the introduction of FTGT, were observed (Figure 1A). These mass additions, which correspond to the monoisotopic mass of one or multiple attached pentose residues, were detected in all three analyzed FTGT-expressing lines (Supplementary Table S2). Characteristic reporter ions of *N*-glycan fragments bound to pentoses were detected on MS2 spectra for all rhEPO glycopeptides (Supplementary Figure S1). This indicates an attachment of the pentoses to the *N*-glycans and not directly to the peptide backbone.

**FIGURE 1.**
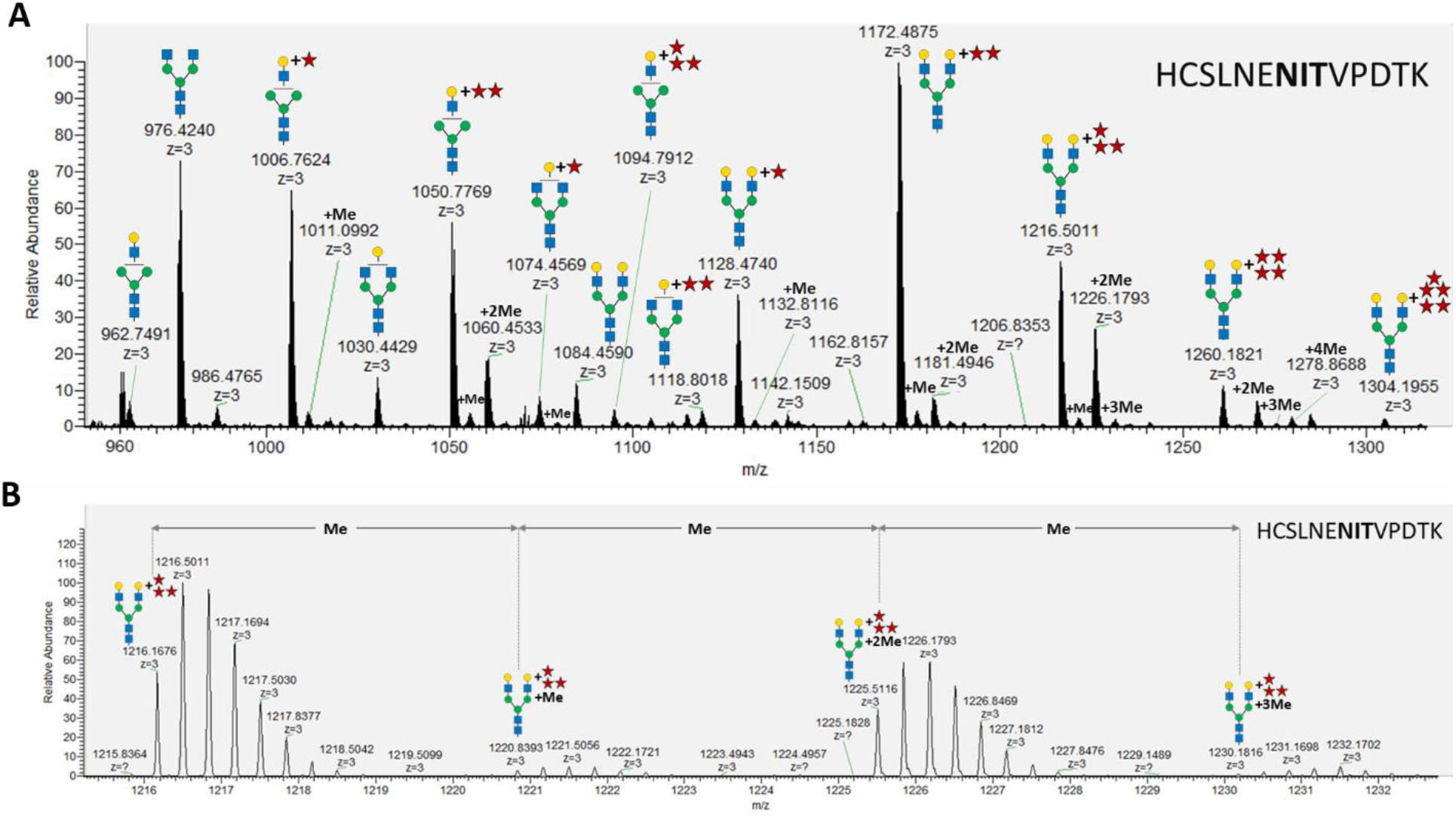
MS analysis of X24-derived rhEPO glycopeptide HCSLNENITVPDTK. **(A)** Precursor mass range (27.6-28.7 min) displaying the X24-derived triple charged ([M+3H^+^]^+^) glycopeptide HCSLNE**NIT**VPDTK with different *N*-glycans attached. **(B)** An average mass shift of 14.0157 Da between the calculated single charged ([M+1H^+^]^+^) precursor masses (from left to right: 1216.1676 ([M+3H^+^]^+^) -> 3646.4882 ([M+1H^+^]^+^); 1220.8393 ([M+3H^+^]^+^) -> 3660.5033 ([M+1H^+^]^+^); 1225.5116 ([M+3H^+^]^+^) -> 3674.5202 ([M+1H^+^]^+^); 1230.1816 ([M+3H^+^]^+^) -> 3688.5302 ([M+1H^+^]^+^)) indicates the addition of methyl groups. Me: methyl, monoisotopic mass shift: +14.0157 Da. *N*-glycans are depicted according to the glycan nomenclature of the Consortium for Functional Glycomics (http://www.functionalglycomics.org), while the attachment of an unknown pentose is depicted as a red star.

While up to three pentoses were detected on mono-antennary galactosylated *N*-glycans, mass shifts corresponding to up to five pentoses were measured on di-antennary galactosylated *N*-glycans (exemplarily depicted in figure 1A for the rhEPO glycopeptide HCSLNE**NIT**VPDTK). Additionally, some pentosylated *N*-glycan structures carried single or multiple mass increments of 14.0157 Da, characteristic for methyl groups. These mass increments occurred as one or up to the number of attached pentoses (Figures 1A, B, Supplementary Figure S2). From this analysis it was not immediately obvious if the detected structures were methyl-pentoses or deoxy-hexoses (e. g. fucoses), as the monoisotopic mass of a deoxy-hexose matches that of a methyl-pentose.

### 3.2 Western blot of rhEPO with an arabinose-specific antibody

As a first step to identify the nature of the unknown pentoses attached to *N*-glycans, proteins recovered from the culture supernatants of the β1,4-galactosylating moss line X24, the parental line 174.16, and line Δ*galt1* (devoid of any *N*-glycan galactosylation), were analyzed via Western blot with the antibody LM6-M, which recognizes short α-L-1,5-arabinan chains (Cornuault et al., 2017). For each line a strong and defined signal at a high molecular weight range (>180 kDa) was observed, which in Physcomitrella is known to be associated with arabinogalactan-proteins (Lee et al., 2005). In the lower molecular weight range, a signal of around 37 kDa was detected exclusively in the X24 sample (Figure 2A). To check if this signal is related to rhEPO, a subsequent anti-hEPO detection was performed (after antibody stripping from the membrane). This anti-hEPO immunodetection revealed rhEPO-corresponding signals between 27 and 37 kDa in all analyzed lines (Figure 2B). The signal with the lowest molecular weight was detected in Δ*galt1*, which displays the most reduced glycosylation pattern of the three investigated lines. The intermediate signal was derived from the line 174.16, while the signal with the highest molecular weight, ranging from 30 to 37 kDa, was detected in X24, which fits with an increased molecular weight of the rhEPO-attached *N*-glycans due to additional galactosylation and pentosylation. This upper part of the rhEPO-corresponding band detected in line X24 overlaps with the position of the signal detected with the LM6-M antibody (Figure 2A). Therefore, we conclude that the LM6-M antibody detects arabinoses on rhEPO produced in moss line X24.

**FIGURE 2.**
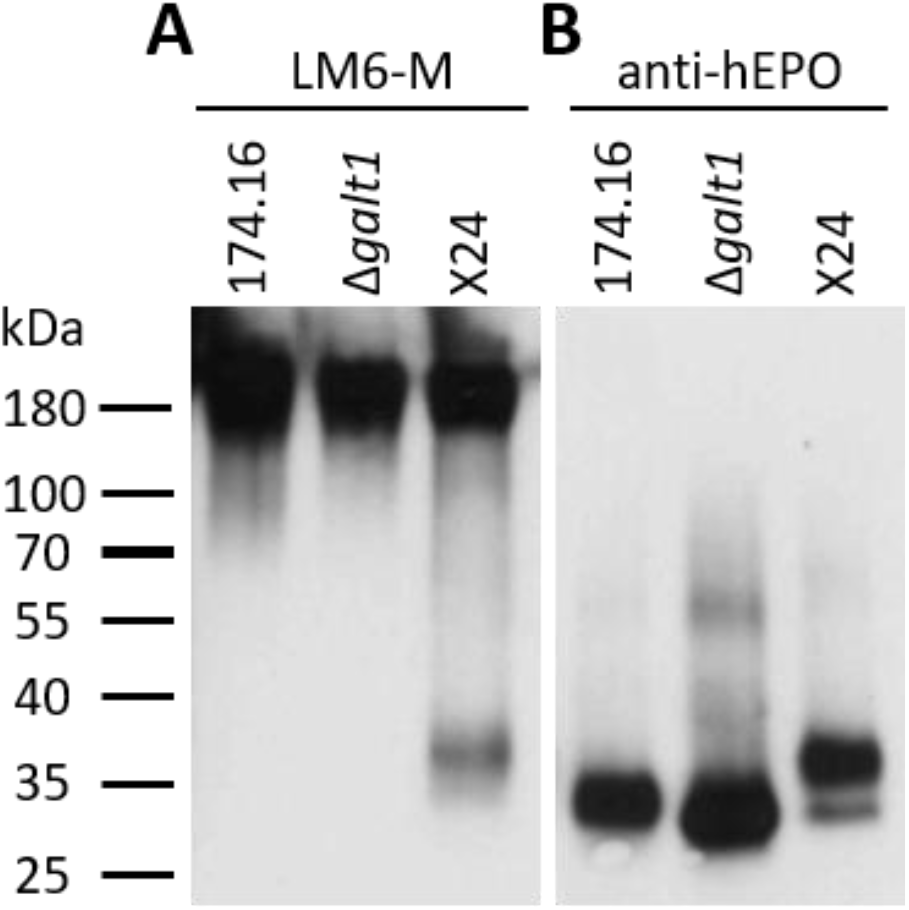
Western blots of precipitated culture supernatants of rhEPO-producing Physcomitrella lines. Five microgram total protein of the precipitated and blotted culture supernatants of the rhEPO-producing lines 174.16, Δ*galt1* and X24 were subsequently immunodetected with the anti 1,5-α-L-arabinan antibody (LM6-M, 1:10) **(A)** and the anti-hEPO monoclonal antibody (1:4,000) **(B)**.

### 3.3 Specific enzymatic activity of α-L-arabinofuranosidase on X24-produced rhEPO confirmed arabinosylation

To further investigate the detected arabinose residues attached to β1,4-galactosylated rhEPO *N*-glycans, samples of all three rhEPO-producing lines were digested with α-L-arabinofuranosidase. The enzyme-treated samples were first analyzed via immunodetection with LM6-M antibodies followed by a detection with anti-hEPO antibodies and compared to mock-treated samples, as a control for possible non-enzymatic hydrolysis.

With the α-L-arabinan-detecting LM6-M antibody, samples treated without α-L-arabinofuranosidase show a similar band profile to the untreated samples analyzed previously. Only in the sample from moss line X24 could a band in the lower molecular weight range of about 37 kDa be detected. Strong LM6-M-derived signals for all undigested samples were observed above 180 kDa, corresponding to arabinogalactan-proteins (Figures 2A and 3A). These high-molecular weight signals disappeared from the α-L-arabinofuranosidase-digested samples, supporting the activity of the enzyme, which is able to digest the 1,5-linked arabinans known to be attached to arabinogalactan-proteins in Physcomitrella (Lee et al., 2005). Furthermore, the arabinose-specific LM6-M-derived signal also disappeared from the digested X24 sample (Figure 3A). The rhEPO-corresponding signals, however, were detected in all samples with the hEPO-specific antibody Western blot (Figure 3B), supporting the hypothesis that the absence of an arabinose-specific signal after α-L-arabinofuranosidase digest is due to the loss of *N*-glycan-attached arabinoses on rhEPO in the β1,4-galactosylating line X24. Moreover, the digestion of the X24-derived sample with α-L-arabinofuranosidase leads to a lower degree of rhEPO-microheterogeneity. This becomes obvious in the comparison of the broad anti-hEPO-derived signal ranging from 27-40 kDa in the undigested sample, which is after α-L-arabinofuranosidase digestion reduced to a lower molecular weight range between 27 kDa and 36 kDa (Figure 3B).

**FIGURE 3.**
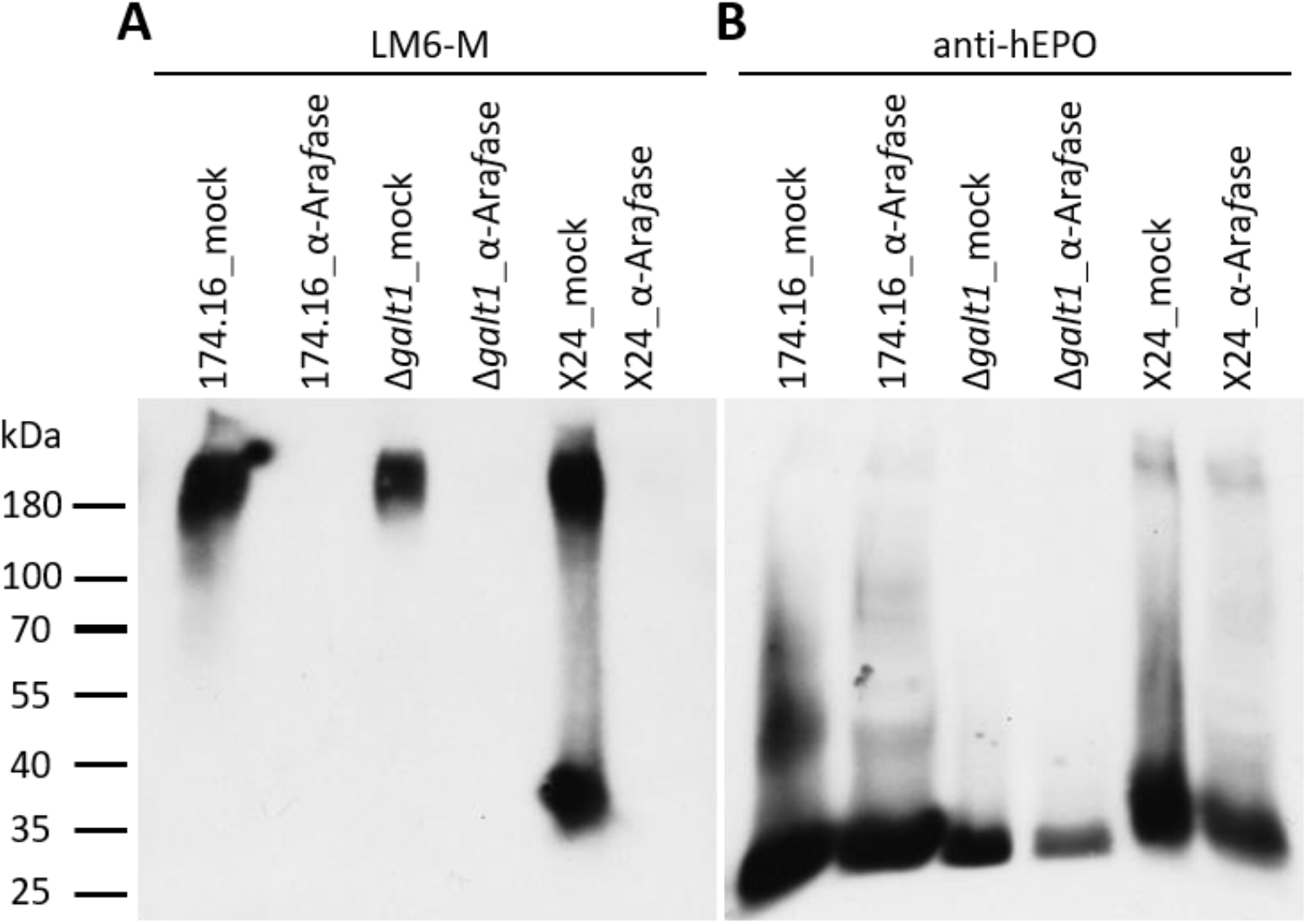
Western blots of mock-treated and α-L-arabinofuranosidase-digested samples from rhEPO-producing Physcomitrella lines. Ten microgram total protein of precipitated culture supernatants of the rhEPO-producing lines 174.16, Δ*galT1* and X24 were digested with one unit of α-L-arabinofuranosidase (α-Ara*f*ase), while control samples were treated equivalently but without α-L-arabinofuranosidase (mock). After separation on SDS-PAGE and blotting, the PVDF-membrane was subsequently incubated with the anti 1,5-α-L-arabinan antibody (LM6-M, 1:10) **(A)** and an anti-hEPO monoclonal antibody (1:4,000) **(B)**.

### 3.4 Mass-spectrometric validation of enzymatic digestion of arabinoses on rhEPO produced in the β1,4-galactosylating moss line

The exact effect of the α-L-arabinofuranosidase treatment on rhEPO-glycopeptides of the β1,4-galactosylating line X24 was further analyzed in triplicates via mass spectrometry in comparison to undigested samples. The total *N*-glycan distribution in rhEPO was estimated by adding together quantitative values (peak areas) from detected glycopeptides. Values were further added together for *N*-glycan classes across all three rhEPO *N*-glycosylation sites (Figure 4). For easier comparison, the quantitative values of all pentose-carrying *N*-glycan structures were also added together, in order to distinguish the total pentosylated and non-pentosylated proportion of an identified structure (Figure 4A). A detailed breakdown of all identified structures and modifications is given in the supplementary table S4 and the quantification is depicted in the supplementary figure S2. The MS data of X24-derived rhEPO glycopeptides from α-L-arabinofuranosidase-digested and undigested samples showed a total amount of galactosylated *N*-glycans of about 66% each. Also the proportion of mono-and di-antennary processed structures within the galactosylated fraction was the same in both conditions, approximately 35% and 65%, respectively. However, in the undigested samples 92% of the galactosylated *N*-glycans were found to be pentosylated, while in the α-L-arabinofuranosidase-treated samples only 29% of the galactosylated structures remained pentosylated (Figure 4A). The number of pentoses on galactosylated *N*-glycans in the undigested approach were 28% single, 31% double, 24% triple, 8.5% quadruple and less than 1% quintuple attachments. In the α-L-arabinofuranosidase treated samples pentoses were evenly cleaved, including a high amount of structures with complete pentose removal, which leads to a clear increase of the corresponding *N*-glycan structure with terminal galactose (Figures 4A, 4B). This suggests that an inefficient digestion is responsible for the attached pentoses detected after α-L-arabinofuranosidase treatments.

**FIGURE 4.**
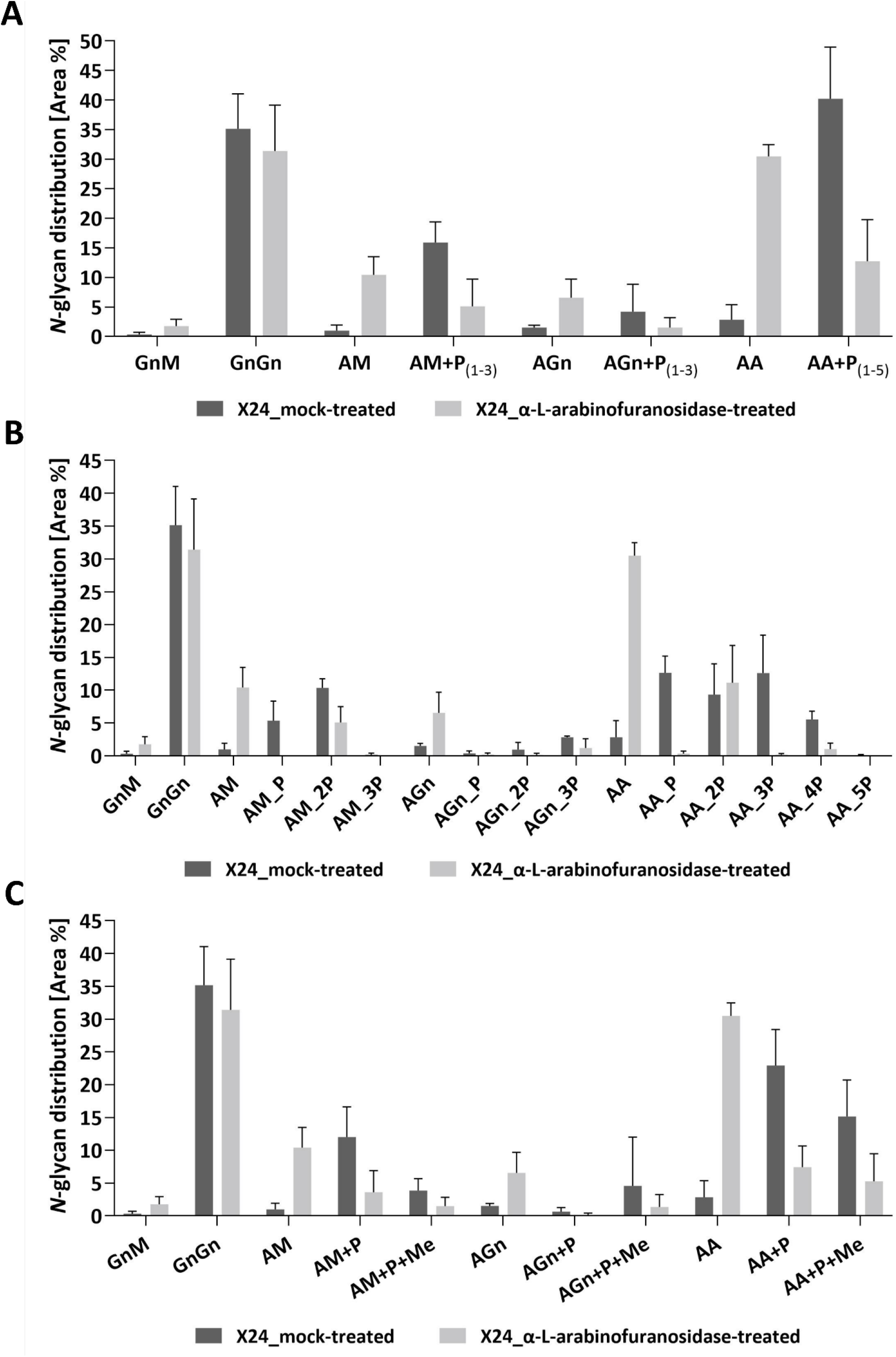
Quantitative MS/MS analysis of the *N*-glycan distribution on rhEPO from α-L-arabinofuranosidase treated in comparison to mock-treated samples. Prior to MS analysis X24-derived rhEPO containing samples were digested with α-L-arabinofuranosidases, and mock-treated samples without enzyme addition were prepared in parallel. Quantitative values are derived from detected glycopeptides **(A).** *N*-glycosylation patterns of rhEPO are represented as relative percentages of all identified *N*-glycan structures within a category (α-L-arabinofuranosidase-treated or mock-treated). For easier comparison, the quantitative values of all pentose-carrying *N*-glycan structures were further added together, thus from each structure the total non-pentosylated and, if applicable, pentosylated shares are depicted. The presence of pentoses on *N*-glycan structures is displayed as +P, while the range of detected pentoses on the corresponding structure is given in subscripted numbers. For a more detailed representation of the data, the pentosylated share of each *N*-glycan structure was further depicted according to the defined number of pentoses (indicated as nP) identified on the respective structure **(B)**. A quantitative profile depicting the share of methylation (+Me) on the identified *N*-glycan structures is given in **(C).** Depicted is the mean of 3 replicates with standard deviation.

Finally, we analyzed the single and multiple mass increments of 14.0157 Da in digested and non-digested samples. The fraction of pentosylated structures with 14.0157 Da increments remained constant after digestion with α-L-arabinofuranosidases. In both treatments about 40% of the pentosylated glycans carried this modification, indicating that the α-L-arabinofuranosidases activity decreased the amount of all pentosylated structures, regardless of the presence of the 14.0157 Da mass increments (Figure 4C).

As the maximal number of identified 14.0157 Da mass additions on pentosylated *N*-glycan structures matches the number of attached pentoses (Figure 1) and these additions did not interfere with the specific arabinose-cleaving activity of the enzymes (Figure 4C), we conclude that the detected masses attached to the galactosylated *N*-glycans are arabinoses, which are occasionally methylated.

## 4 DISCUSSION

Glycosylation, a frequent and complex posttranslational modification of proteins, is a critical quality feature for glycoprotein-based therapeutics, as it influences their conformation, solubility, activity, pharmacokinetics and antigenicity (Arnold et al., 2007; Solá and Griebenow, 2010). The composition of the respective *N*-glycans is dictated by intrinsic characteristics of the protein itself, such as conformation, as well as by the glycan-processing enzymes of the production platform (Clausen et al., 2015; Suga et al., 2018). *N*-glycosylation of most biopharmaceutical production hosts, even the predominantly used mammalian cell systems, differ to their human counterparts to different extents (Wang et al., 2015). For instance, *N*-glycolylneuraminic acid (Neu5Gc), a sialic acid not existing in humans and consequently associated with antibody formation (Tangvoranuntakul et al., 2003; Padler-Karavani et al., 2011), can be found on *N*-glycans of glycoproteins produced in some non-human mammalian cell lines (Varki, 2001; Ghaderi et al., 2012). Although plant *N*-glycosylation differs from the human pattern, its humanization, which includes the removal of plant-specific sugar residues, the introduction of a β1,4-galactosylation capacity and the final establishment of terminal *N*-glycan sialylation, has been performed to varying degrees in different plant systems reviewed in Montero-Morales and Steinkellner (2018). These studies have demonstrated a great flexibility of plants towards glyco-engineering. Especially the moss Physcomitrella offers the additional advantages of a high rate of homologous recombination in mitotic cells, a characteristic feature used for efficient precise genome editing, and a haploid gametophytic tissue, enables immediate implementation of glyco-modifications(Parsons et al., 2012; Decker et al., 2014; Wiedemann et al., 2018).

The β1,4-linked galactoses on *N*-glycans provide the anchor for sialic acid, but terminal galactose also plays an important role in non-sialylated glycoproteins. For example, asialo-EPO was proposed to be neuroprotective (Erbayraktar et al., 2003; Peng et al., 2020) and on the Fc domains of monoclonal antibodies terminal *N*-glycan galactosylation increases complement-dependent (Hodoniczky et al., 2005) as well as antibody-dependent cytotoxicity (Thomann et al., 2016).

In terms of *de novo* β1,4-galactosylation in plants, its efficiency and degree of maturation (mono- or di-antennary) depends on the expression level of the respective galactosyltransferase (Kallolimath et al., 2018), its Golgi localization, determined by the N-terminal CTS domain (Strasser et al., 2009; Hesselink et al., 2014; Kriechbaum et al., 2020), and the reporter glycoprotein itself (Schneider et al., 2015; Kriechbaum et al., 2020).

In this study, we established β1,4-galactosylation on rhEPO produced in moss devoid of plant-specific sugar residues. To target the β1,4-GalT activity to the late Golgi compartments, the catalytic domain of this enzyme was fused to the CTS domain of the moss-endogenous α1,4-fucosyltransferase, whose activity is the last known in plant *N*-glycan maturation (Fitchette et al., 1999; Parsons et al., 2012).

Stable expression of this chimeric variant, FTGT (Bohlender et al., 2020), in an rhEPO-producing moss line resulted in 66% rhEPO galactosylation. 65% of all galactosylated *N*-glycans were mature di-antennary processed ones, indicating a medial- to trans-Golgi localization of the FTGT enzyme. These values are very promising, considering that previous studies reported lower galactosylation efficiencies with high degrees of mono-antennary galactosylation on rhEPO produced in *N. benthamiana* or *N. tabacum* plants (Kittur et al., 2013; Kriechbaum et al., 2020). However, the galactosylation efficiency on rhEPO produced in *N. benthamiana* was increased by knocking out the β-galactosidase NbBGAL1, an enzyme responsible for galactose cleavage (Kriechbaum et al., 2020). A similar strategy might be applied to moss.

Accompanying the established human-like galactosylation, we detected the attachment of pentose residues on β1,4-galactosylated *N*-glycans. Up to three pentoses were attached to mono-antennary and up to five pentose residues to di-antennary galactosylated *N*-glycans, which indicates the building of short pentose chains. These were not present in the corresponding parental line with an intact β1,3-galactosyltransferase (Parsons et al., 2012), indicating that naturally occurring β1,3-galactosylated *N*-glycans do not display a substrate for this modification.

*In planta N*-glycan pentosylation on a recombinant protein upon the establishment of β1,4-galactosylation has also been observed in *N. tabacum* (Kittur et al., 2020), suggesting that this phenomenon is not restricted to Physcomitrella but rather affects plant-based production in general. Pentosylation was also observed in sialylating moss lines (Bohlender et al., 2020). However, in these plants either pentoses or sialic acid could be detected on galactosylated *N*-glycans, indicating that the pentosylation may interfere with the full *N*-glycan humanization of plant-derived glycoproteins. This observation confers importance to the elucidation of the respective pentose residues.

Based on immunodetection with LM6-M, a monoclonal antibody recognizing short chains of α1,5-linked arabinan (Cornuault et al., 2017), we could identify the pentoses on moss-produced rhEPO as arabinoses. Specific digestion of these pentoses with α-L-arabinofuranosidase, an enzyme specifically cleaving α1,2-, α1,3- and α1,5-linked arabinoses from arabinan molecules, was verified via immunodetection supported by MS analysis of rhEPO glycopetides. These findings confirm the identity of the pentoses as (short chains of) α-linked arabinoses. Additionally, we found the arabinoses to be occasionally methylated. Some residual pentoses after α-L-arabinofuranosidase digest may be attributed to poor hydrolysis of α-1,5-linked arabino-oligosaccharides by the enzymes used.

Recently, the presence of arabinose and methylated arabinose on *N*-glycans of the microalga *Chlorella sorokiniana* has been described (Mócsai et al., 2020). However, this sugar has never been observed on *N*-glycans of Physcomitrella before the establishment of human-like β1,4-galactosylation. Evidently, an arabinosyltransferase from a different biosynthetic pathway recognizes the β1,4-galactosylated *N*-glycan as substrate. Plants display a wide diversity of cell-wall glycans and O-glycosylated hydroxyproline-rich glycoproteins (Seifert et al., 2021). This diversity originates from the combination of different monosaccharides and various linkages, generated by a huge variety of glycosyltransferases from which a considerable amount has not been thoroughly characterized yet (Showalter and Basu, 2016; Amos and Mohnen, 2019). Some enzymes responsible for the attachment of arabinoses to β1,4-linked galactoses on O-glycosylated arabinogalactan proteins as well as in cell-wall associated structures like rhamnogalaturan I have been described, but many still remain unknown (Léonard et al., 2010; Laursen et al., 2018; Ropartz and Ralet, 2020; Petersen et al., 2021). The identification of the enzyme or enzymes responsible for the arabinosylation of galactosylated *N*-glycans is therefore not a straightforward task.

For the application of plant-based biopharmaceuticals, this newly appearing *N*-glycan attachment bears the risk of immunogenicity in patients, as arabinose is a sugar not produced in humans (Anderson et al., 1984; Steffan et al., 1995; Leonard et al., 2005). To avoid arabinose attachment, the arabinosyltransferase activity might be bypassed by a chimeric β1,4-galactosyltransferase acting very late in the trans-Golgi apparatus. Alternatively, the responsible arabinosyltransferases need to be identified and abolished by gene targeting to create stable lines devoid of *N*-glycan arabinosylation. To this aim our study provides the first important step by elucidating the unknown pentose residues, which helps to ensure the production of safe biopharmaceuticals in plant-based systems.

## 5 Data availability

All data generated in this study is included in this paper and the supplementary information. The mass spectrometry proteomics data have been deposited to the ProteomeXchange Consortium via the PRIDE partner repository with dataset identifier PXD030443.

## 6 Author Contributions

LLB performed most of the experiments, SNWH performed the MS data analysis, NB performed some Western blot experiments, FRJ created the analyzed lines I10, X13 and X24, LLB, JP, RR and ELD designed the study and wrote the manuscript.

## 8 Acknowledgements

We thank Agnes Novakovic for technical support to this work. We thank Prof. Dr. Bettina Warscheid for the use of the QExactive Plus instrument, Prof. Dr. Jörn Dengjel and Dr. Verónica I. Dumit for the use of the QTOF instrument and Anne Katrin Prowse for proof-reading of the manuscript.

## Supplementary Figures

**SUPPLEMENTARY FIGURE S1.**
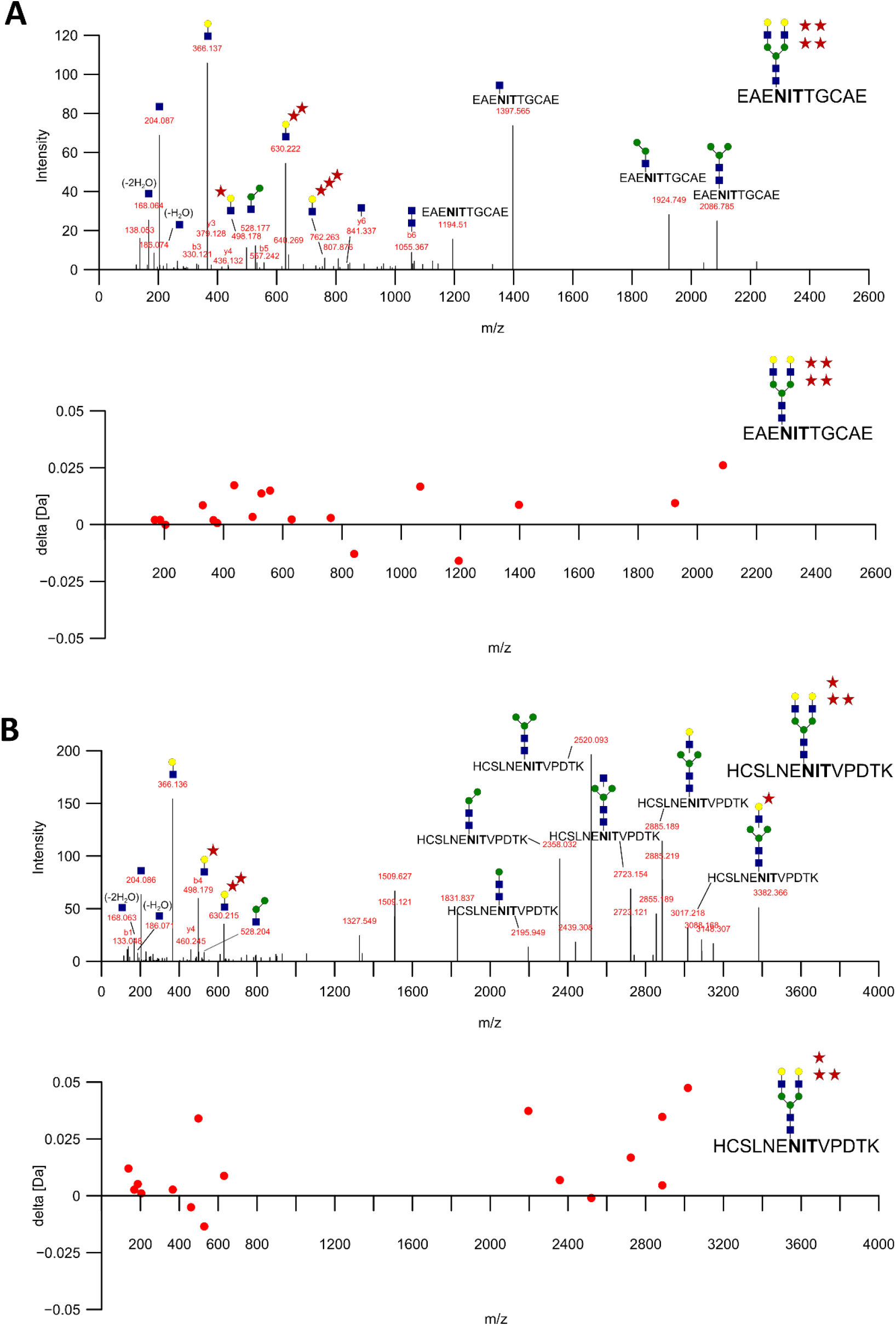
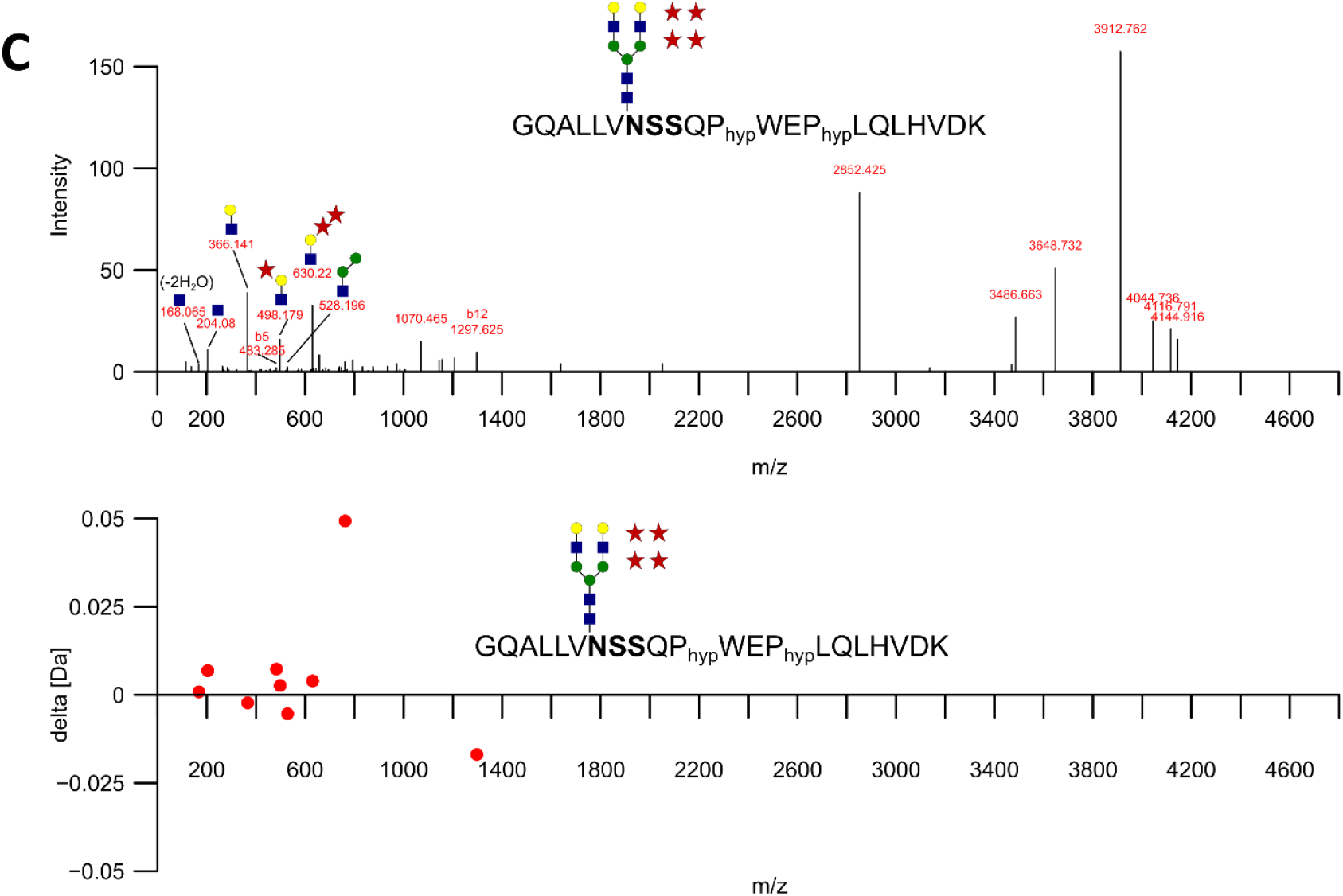
Collision-induced dissociation (CID) fragment spectra of di-antennary galactosylated and additionally pentosylated glycopeptides representing the three rhEPO *N*-glycosylation sites (Asn24, Asn38, Asn83). **(A)** CID fragment spectrum of the identified glycopeptide EAE**NIT**TGCAE ([M^+^3H^+^]^+^ = 1115.7574). The identified precursor mass corresponds to a di-antennary galactosylated *N*-glycan with four attached pentoses. **(B)** CID fragment spectrum of the identified glycopeptide HCSLNE**NIT**VPDTK ([M^+^4H^+^]^+^ = 912.3763). The identified precursor mass corresponds to a di-antennary galactosylated *N*-glycan with three attached pentoses. **(C)** CID fragment spectrum of the identified glycopeptide GQALLV**NSS**QPhypWEPhypLQLHVDK ([M^+^4H^+^]^+^ = 1136.2589). The identified precursor mass corresponds to a di-antennary galactosylated *N*-glycan with four attached pentoses. The mass shift of carbamidomethylation (+57.0214 Da) is added to all contained cysteine residues. Hydroxylation of a proline is indicated by Phyp (+15.9949 Da). Spectra were acquired on a Q-TOF instrument from a sample of line X24. The monoisotopic masses of the detected sugar reporter ions are as followed: [GlcNAc]^+^ (blue square) = 204.0867, [GlcNAc - H_2_O]^+^ = 186.0761, [GlcNAc - 2H_2_O]^+^ = 168.0655, [GlcNAcHex]^+^ = 366.1395, [GlcNAcHex_2_]^+^ = 528.1923, [GlcNAcHexPent]^+^ = 498.1818, [GlcNAcHexPent_2_]^+^ = 630.2241; [GlcNAcHexPent_3_]^+^ = 762.2664 with Hex= Hexose = galactose (yellow circle) or mannose (green circle) and Pent = pentose (red star). *N*-glycosylation consensus sequences of depicted glycopeptides are shown in bold. Below each fragment spectrum the distribution of the mass errors of identified fragments is shown.

**SUPPLEMENTARY FIGURE S2.**
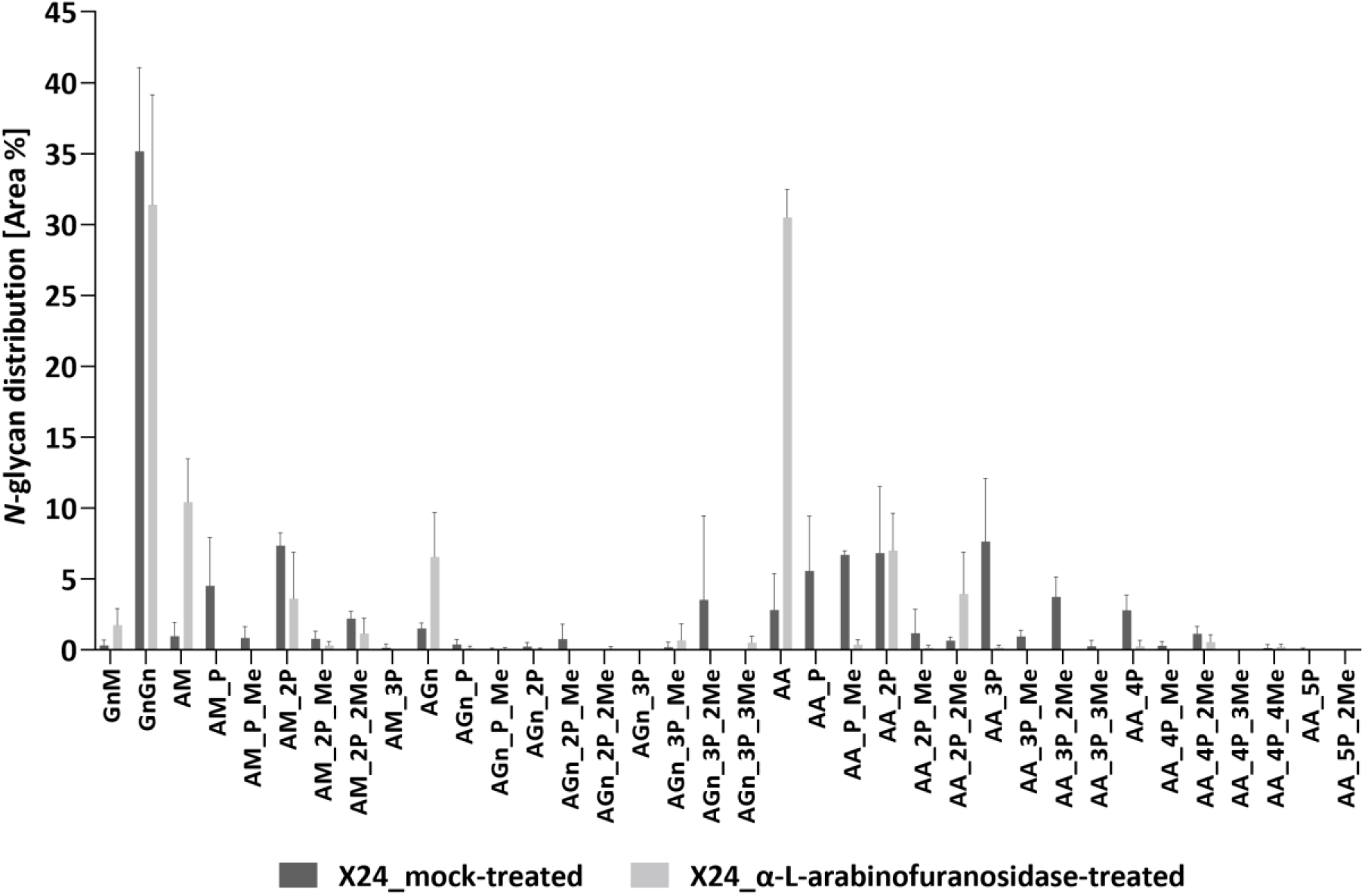
Quantitative MS/MS analysis of the N-glycan distribution on rhEPO from α-L-arabinofuranosidase treated vs. mock-treated samples. Prior to MS analysis rhEPO-containing samples of line X24 were digested with α-L-arabinofuranosidase, while mock-treated samples without enzyme addition were prepared in parallel. For MS analysis trypsin and GluC-released rhEPO glycopeptides were analyzed. The *N*-glycosylation pattern of α-L-arabinofuranosidase-treated and mock-treated glycopeptides is represented as relative percentages of all identified *N*-glycan structures across all three *N*-glycosylation sites. P: Pentose, Me: Mass increment of 14.0157 Da corresponding to methylation. The mean of three replicates with standard deviation is depicted.

